# AG3D: A low-cost educational 3D printable toolkit for agarose gel electrophoresis

**DOI:** 10.1101/2023.03.22.533785

**Authors:** Matthew W Barwick, Sidne Fanucci, Siso Mbanxa, Khanyisile Buthelezi, Michael D Jukes, Garth L Abrahams, Earl Prinsloo

**Affiliations:** Biotechnology Innovation Centre, Rhodes University, P.O. Box 94, Makhanda, 6140, South Africa; Department of Biochemistry & Microbiology, P.O. Box 94, Rhodes University, Makhanda, 6140, South Africa

**Keywords:** 3D Printing, agarose gel electrophoresis, low-cost, DNA separation

## Abstract

Access to desktop additive manufacturing is growing and the argument could be made for 3D printers to be standard laboratory equipment. The power of the printers lies in the democratisation of scientific equipment. Traditional agarose gel electrophoresis forms a cornerstone of molecular biology research, teaching and learning. Reliable electrophoresis is dependent on a number of factors which include standardized commercial equipment with respect to casting trays, combs, horizontal tanks and power supplies. The aforementioned systems come at a high initial cost; this is before factoring in the costs of standard electrophoresis grade agarose and associated reagent pricing. Here, we present a basic method for the additive manufacturing of a simple 3D printable agarose gel electrophoresis (AGE) unit with a built-in gel casting tray for standard size microscope slide-based AGE; named AG3D. The system presented was validated using different standard agarose-buffers (Tris Acetate EDTA and Tris Borate EDTA) and commercially available base-pair ladders. We provide a comparison between the AG3D and a commercial AGE system in respect to resolving power of the electrophoresis unit and discuss the overall reduction in cost afforded by the AG3D electrophoresis toolkit. The method presented has the potential for application in low resource educational environments by:

Significantly lowering costs through the reduction of reagents (agarose, buffers etc).
Allow for the use of low sample volumes.
Providing an open-source toolkit for modification whether for research or teaching & learning

**Graphical abstract:** 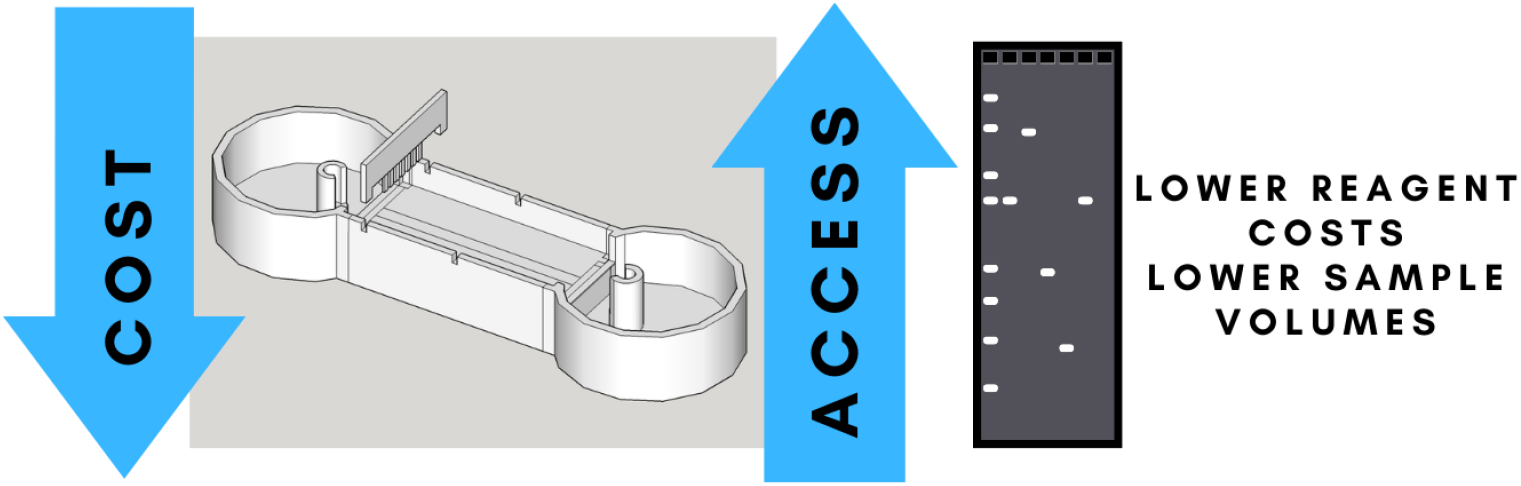

**Specifications table:** 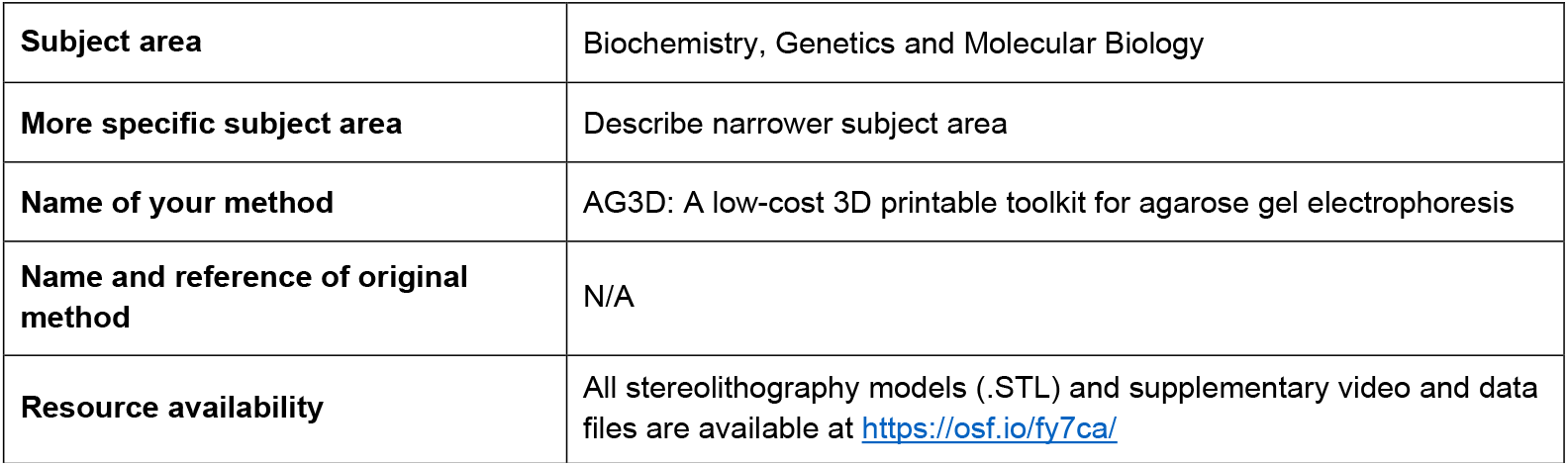

## Method details

### Background

Ethidium bromide agarose gel electrophoresis (EtAGE) is a standard technique in any molecular biology laboratory investigation that allows for efficient separation of molecular DNA leading to sizeable and identifiable segments; a technique that has remained largely untouched and follows the basic principle as outlined by Aaij and Borst [1]. Commercially available gel tanks with buffer reservoirs housing platinum wires as electrodes are costly and sometimes beyond the reach of some laboratories. This is compounded by the reagent costs e.g. DNAse/RNAse-free electrophoresis grade agarose and the individual buffer components adding to the overall cost of the electrophoresis system. The high cost of these systems provides a need for easily accessible, rapid, and lower cost systems [2] in order to erase the barrier to entry for even the most basic molecular biology laboratories and teaching environments.

Three-dimensional (3D) printing of lab equipment has become popular in recent years due to the ability to design, rapidly prototype and produce low-cost equipment for laboratory use. One of the most common technologies of 3D printing is fused deposition modelling/fused filament fabrication (FDM/FFF) which translates the computer-aided design (CAD) model by extruding thermoplastic filament layer by layer in X, Y and Z dimensions to create the 3D structure by deposition through a heated nozzle. There is a wide variety of materials to choose from when it comes to 3D printing, but ultimately the choice is dependent on the 3D printer used, the function of the proposed equipment, and the available funds within the research laboratory. Acrylonitrile butadiene styrene (ABS) and Polylactic acid (PLA) are two of the most widely accessible and low-cost 3D printing polymers [3]. Three-dimensional printing has rapidly grown in the past decade with the decline in cost of the method, the technology allows for designing, prototyping, and manufacturing of simple to complex objects, compared to traditional manufacturing process, 3D printing allows for rapid and cost-effective manufacturing. Due to the cost-effective benefit, many research groups have incorporated 3D printers into their laboratories to solve issues in a creative and cost-effective manner items such as: falcon tube racks; microscope phone holders; 96 well plates; to gel combs have been printed by various laboratories [4].

Multiple attempts have been made to modify the EtAGE system for varying reasons. Low-cost solutions have been proposed in Bets & Berard [5] wherein the system was made out of a flexible, plastic and hinged pipette box with electrodes made out of aluminium foil. Ens et al. [6] proposed running an electrophoresis system using common items found in a house – this included using household plastic containers, utilising folded aluminium foil to create electrode strips, and using five 9V batteries in series were used as the power supply. Adamski et al. [7] used additive manufacturing to develop a lab-on-chip for single sample DNA separation using a 3D printed microfluidic system for efficient separation of DNA ranges from 50-800 bp in 10 min in 200 V/cm.

The focus of this study was to design a simple reusable miniature electrophoresis toolkit (Fig 1) that significantly reduces the costs of routine agarose gel electrophoresis. The system was designed to fit a standard 20 × 20 cm Fused Deposition Modelling/Fused Filament Fabrication (FDM/FFF) print bed and be manufactured as a single unit without the use of support materials. Further cost reductions were envisioned through the use of low-cost copper single wire connectors as electrodes and reagent quantities (one tenth standard volumes). A modification which utilises standard platinum wire as electrodes was also designed. The system presented here, should allow for routine use in research and teaching environments.

**Fig 1.**
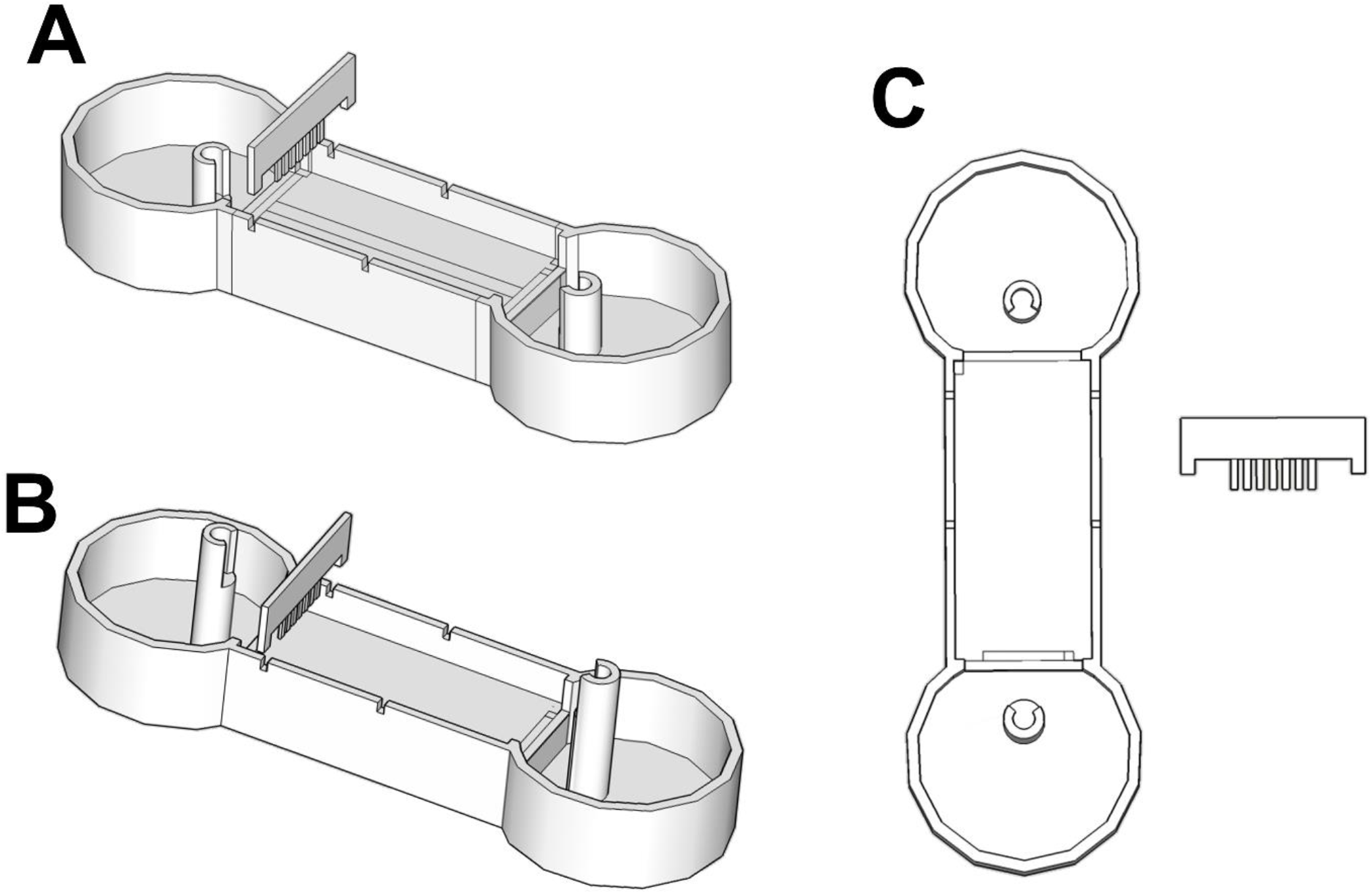
Schematic illustration of the AG3D electrophoresis tanks and combs. Rendering of the .STL file prepared in Trimble SketchUp (v 17.2.2555). A – Side view of schematic AG3D model using banana plugs only. B – Side view of the second schematic AG3D model using banana plugs with the addition of platinum wire. C – Top view of schematic AG3D model.

### Fabrication of the AG3D electrophoresis system

The AG3D electrophoresis tanks were designed using Trimble SketchUp (v 17.2.2555) and printed using relatively low-cost fused filament fabricator (FFF) 3D printers; a cartesian Prusa i3 clone Wanhao Duplicator i3 v2.1 (3D Printing Store Pty LTD, South Africa) and an Kossel Delta based AnyCubic® Kossel Linear Plus 3D printer (DIYElectronics, South Africa). All printers were equipped with 0.4 mm sized print nozzles. The tanks and gel combs were printed in 1.75 mm diameter polylactic acid (PLA, Verbatim or ESUN PLA+) at 100 µm layer height with a solid infill (105% filament flow) to ensure water tightness. Two variants were designed based on the electrodes to be used i.e. standard copper plated 39 mm multimeter banana plugs (single wire connectors) or in combination with standard platinum (0.2 mm x 125 mm) wire. The latter was designed to ensure that only the platinum electrodes were submerged to avoid galvanic corrosion associated with copper electrodes. The multimeter banana plugs were disassembled to expose the entire connector (Fig 2). All electrodes were spaced to allow for a linear electrical field. A 7 well gel comb (0.75 mm) was designed allowing for the casting of approximately 4 mm thick agarose gels; sample well volumes were calculated to hold 4 µL. The tank reservoir was designed to hold approximately 50 mL of buffer for efficient electrophoresis, therefore approximately 10:1 buffer to gel ratio. The outer walls of the units were covered in a thin layer of Marine Clear Silicone Gel (Bostik, South Africa) to ensure watertightness. Manifold printable models (STL format) are available as supplemental files (https://osf.io/fy7ca/).

**Fig 2.**
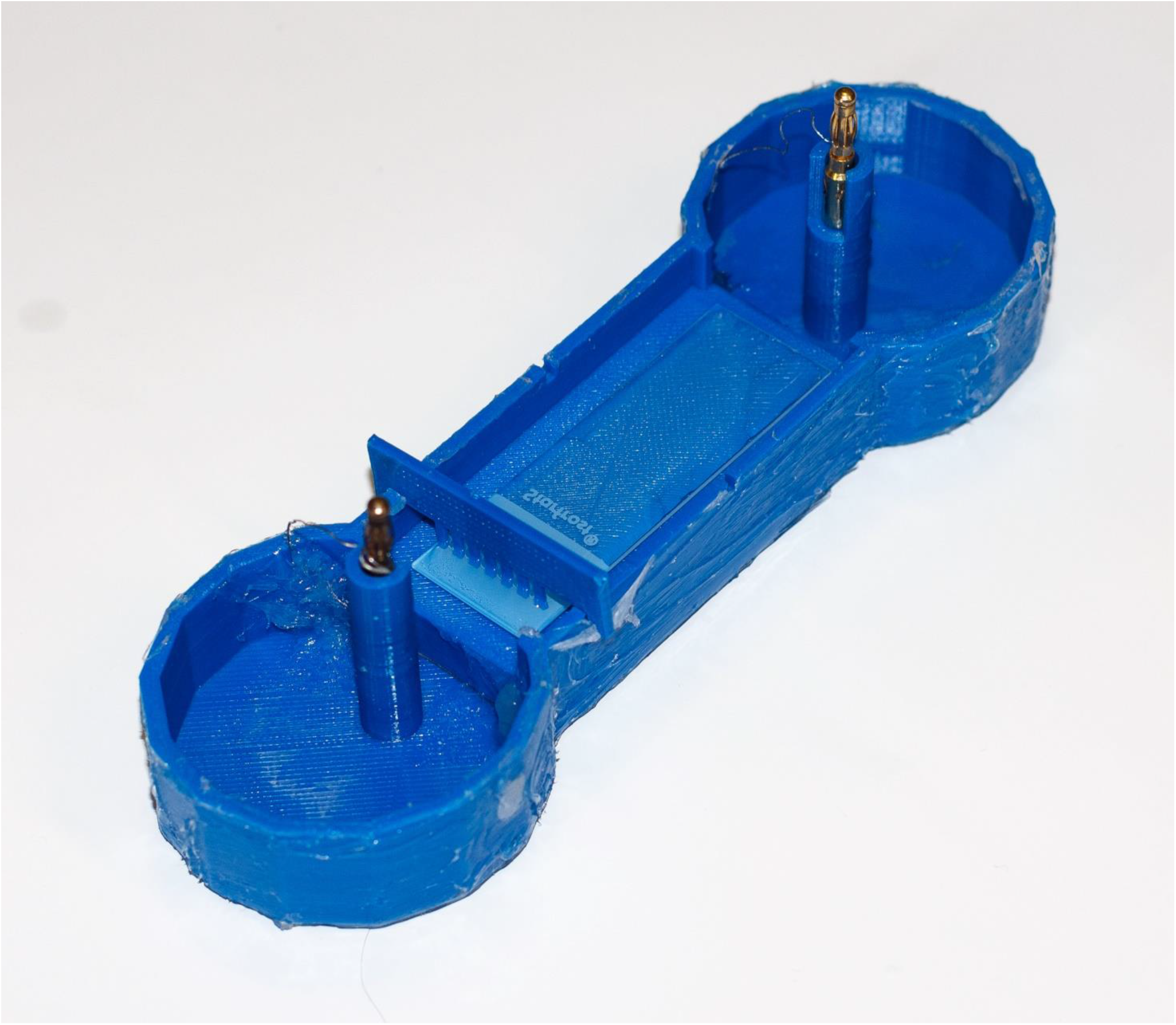
Assembled 3D printed construct of the AG3D system. Image was captured using a Canon 650D camera using a 50mm f1.8mm prime lens.

### Comparative electrophoresis using the AG3D system

Tris-acetate EDTA (TAE) and Tris-borate EDTA (TBE) agarose gels were run on the platinum-electrode, & copper electrode AG3D and 50 mL standard mini agarose gel electrophoresis systems. Ten times strength (10 x) TAE (pH 7.6) and TBE (pH 8.0) buffers were formulated & autoclaved; 1 % (w/v) agarose gels (Seachem LE, Lonza) were prepared using the respective buffers at 1 x concentration. Gels were prepared using a standard 900 W microwave in heat resistant borosilicate Schott bottles containing no ethidium bromide. For the AG3D electrophoresis system, 4 ml of the agarose was aliquoted slowly via 1 ml autopipette on a clean 26 × 76 mm microscope slide (Waldemar Knittel, Lasec) (Supplementary Video 1 available at https://osf.io/fy7ca/). The standard horizontal mini agarose gels were prepared as above, except that 30 mL of agarose solution was used for casting and 250 mL buffer was used to fill the tank.

Two microlitres (2 µL) of 100 ng/μL KAPA Universal DNA ladder (Roche) was used as the sample to compare the different electrophoresis systems and the efficiency of band separation. All gels were electrophoresed for 30 minutes at constant voltage of 100 V (60 mA) using an ENDURO 300W Power Supply (Whitehead Scientific, South Africa). All gels were stained post electrophoresis in ethidium bromide solution (0.5 µM/mL) on a Blotboy lab rocker (Benchmark Scientific) for 30 minutes at room temperature and visualised using the BioRAD ChemiDoc™ XRS+ system. Gel images were captured using Image Lab Software v6.0.1.

## Method Validation

### Printing, Assembly & Robustness of AG3D

Following CAD design finalisation, the AG3D system was printed in PLA. The printing parameters were set at: Infill density, 100%; Extrusion temperature, 220 °C; Raster Angle, slightly below 90°; and Layer Thickness, 0.1 mm. Fig. 2 indicates the printed AG3D Agarose Electrophoresis System coated in silicon following printing and assembly; the variant with a raised copper electrode combined with a more robust platinum wire is shown. Banana plugs were disassembled as described in methodology and inserted in the standoffs designed. The unit assembly for the platinum wire variant is through feeding platinum wire through the banana plug through-hole and fixing the platinum wire electrodes at the cathode and anode using hot glue. The banana plugs are then seated in the appropriate anode and cathode standoff.

A study performed by Fernandes et al., 2018 analysed the combined influence of 3D printing parameters on the mechanical properties of Polylactic acid (PLA) plastic. Four 3D parameters (infill density, extrusion temperature, raster angle and layer thickness) were analysed at three different levels by printing cubes with different 3D printing parameters. The study tested infill density at: 20 %, 40 %, and 60 % and performed mechanical tests that evaluate different tensile tests (Ultimate tensile stress, Yield Strength, Modulus of Elasticity, Elongation at Break, and Toughness). The study found that printing parameters (Infill Density, 60 %; Extrusion Temperature, 220 °C; Raster Angle, 0°/90°; and Layer Thickness, 0.1 mm) indicated the best results for ultimate tensile stress and elongation. The AG3D system has the same set parameters for the extrusion temperature and layer thickness with the parameters mentioned above by Fernandes et al., 2018. The highest infill density evaluated by Fernandes et al., 2018 was 60 % compared with the AG3D system, the AG3D has a higher infill density (100%). Fernandes et al., 2018 found that the higher the infill density, the higher the amount of stress the object can withstand. The decision to print the AG3D as a solid infill was to ensure watertightness.

### Water absorption of AG3D

Jordá-Vilaplna et al., 2014 measured water-contact angle of 3D printed PLA and found that PLA behaves in a hydrophobic manner when interacting with water but noted that the water-contact angle decreased with a decline in surface roughness. Water absorption of 3D printed objects is heavily influenced by porosity, which, in turn, is affected by layer thickness; as layer thickness increases, the number of pores created within each layer of plastic deposited increases, therefore increasing overall porosity [8]. Various studies have demonstrated that lower layer thickness; 0.1 mm, results in more efficient adhesion between layers (i.e. increases compactness of layers) and therefore contributes to a rougher surface which in turn results in lower observed porosity and water absorption [8–10]. Furthermore, the extrusion temperature of PLA also contributes to the level of water absorption of the printed construct. The higher the extrusion temperature, the higher the water absorption. Temperature affects the structure of the polymer by forming microcracks which increases porosity and therefore water absorption. Fernandes et al., 2018; Ayrilmins et al., 2019; and Vincente et al., 2019 assessed the effect of temperature on water absorption of PLA. The same trend was noted whereby the 220°C extrusion temperature resulted in higher water absorption, despite the higher surface roughness, when compared to an extrusion temperature of 200°C., To further mitigate the potential increase in porosity of the system and aid in overall waterproofing, the final 3D printed AG3D was coated with 1-2 layers of silicone.

### Agarose Gel Casting

#### Comparison to Commercial Electrophoresis Systems

Tris-Acetate EDTA (TAE) and TBE agarose gels were used for the two models of the AG3D electrophoresis system, as well as the commercial electrophoresis system and band separation comparisons shown in Fig 3.

**Fig 3.**
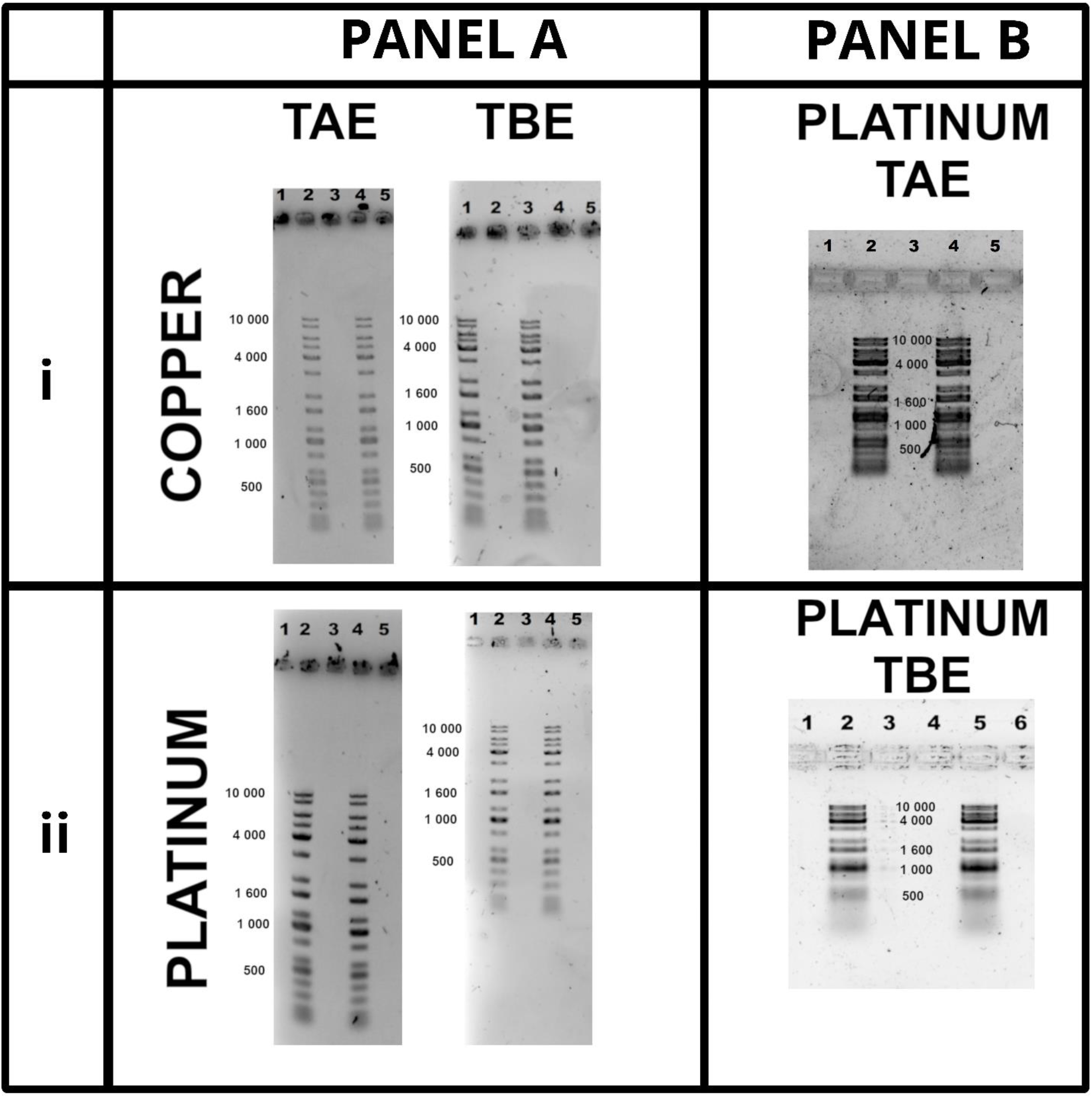
Comparison of band separation between AG3D model with banana plugs/copper electrodes (panel A, i), AG3D platinum wire electrodes (panel A, ii), and commercial mini-gel electrophoresis system (panel B) in TAE and TBE. All gels contained 1% (w/v) agarose with KAPA Universal DNA ladder used as the sample for each gel.

The TAE and TBE copper and platinum electrode variant AG3D agarose gels showed identical band separation, highlighting that the addition of the platinum wire to the AG3D system, had no effect on band separation when compared to the copper banana plug AG3D model, Fig 3, panel A.

When comparing the performance of the AG3D models to the commercial mini agarose electrophoresis system (Fig 3, panel B) there was a distinct difference in band separation when all conditions were kept constant. The AG3D models showed distinguishable band separation and greater band resolution. Bands below 500 bp are visible with the AG3D system, while band resolution declined for bands smaller than 1000 bp when using the commercial system. The AG3D system obtained better band separation in a shorter amount of time, while the commercial system produced more compact bands; this was validated by the standard curves (DNA size versus relative fronts – Supplementary Data File 1 available at https://osf.io/fy7ca/). Standard curves were created for all the gels, indicating the band separation. The AG3D system was able to adequately separate the various DNA fragments with differences noted in the band separation with the use of TBE and TAE. When comparing the different buffers, TBE and TAE, with both the platinum wire and without, the TAE buffer showed more effective band separation than the TBE buffer. Distortion in the migration of the bands with the non-platinum system is noted when compared to the platinum AG3D system. This is likely as a result of the shape of the electric field created by the parallel platinum wires as opposed to the banana plugs.

The DNA separation observed for the commercial system exhibited shorter migration distances, resulting in poor band separation (Fig 3, panel B). Sanderson et al., 2014 noted that DNA electrophoresis is usually performed at 100-150V for 50- 90 mins to discern adequate band separation. The commercial system likely required more time to achieve good electrophoretic separation. While indirect, the time period required for complete electrophoresis using the commercial system could affect overall cost (time, reagents and equipment) of the experiment The buffers (TAE and TBE) used are most commonly used for DNA size determination although they perform similarly, TAE has a better band separation for larger sized DNA while TBE has a better band separation for smaller- sized DNA [11] - [13]. Fig 3 shows that there is little difference between TAE and TBE for both the AG3D and commercial system and the choice of either using TAE or TBE buffer will, therefore, be dependent on the size of DNA undergoing electrophoresis.

Copper was selected as the electrodes for one variant of the AG3D system, the other variant made use of platinum wire as the electrode. Copper has a higher conductivity, lower resistivity, and in terms of costs – copper is cheaper compared to platinum [13]. The longevity of the AG3D system is affected by the durability of the copper electrodes, due to copper undergoing galvanic corrosion during electrophoresis (Fig 4). The corrosion could, however, be mitigated by polishing and reversing the anode and cathode electrodes in alternate electrophoretic experiments.

**Fig 4.**
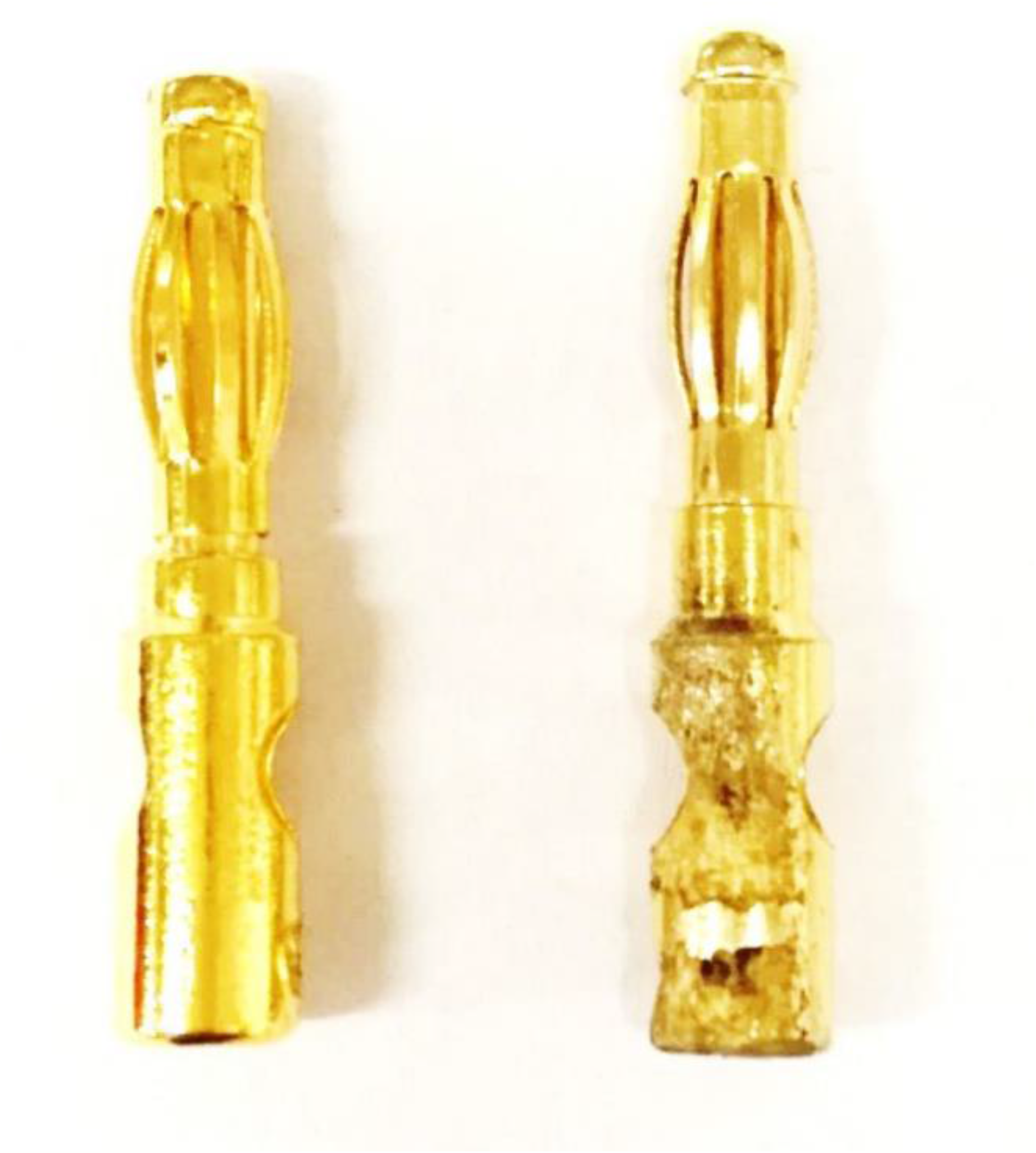
Electrode Surface Corrosion following multiple electrophoretic runs under standard electrophoresis conditions.

Agarose gel electrophoresis is a standard technique in any molecular biology laboratory investigation; a technique that has remained largely untouched and all follow the basic principle as outlined by Aaij and Borst [1]. The system proposed here, utilised the same principle but as a cost reduction exercise to exemplify the power of desktop additive manufacturing in laboratories. The AG3D unit is a simple, compact, low-cost system using copper electrodes, when compared to commercially available gel tanks with platinum wire as electrodes. A platinum wire electrode variant was also presented, which while increasing the cost of the unit does contribute substantially to its longevity. The current working iteration allows for efficient separation of electrophoretic standards. Resolution issues observed in the 100 – 200 bp ranges are most likely as a result of the gel percentage. Surface corrosion on the copper electrode, typically at the anode, can decrease electrophoretic efficiency followed repeated electrophoresis. A proposed solution involving electrode polishing and reversal does ensure increased usability of the lower cost copper electrodes. All considered, the system presented here offers substantial cost reduction even when factoring in the cost of a low-cost desktop 3D printer. Reliable low-cost desktop 3D printers with large enough printbeds can be purchased between $150-$200 and 1 kg of PLA filament averages ∼$24; prices that are comparable and even lower than that of a standard horizontal agarose gel tank. Given the average printed mass of 95 g, the estimated material cost per tank unit is $2.28 combined with the average cost of a pair of banana plug connectors (∼$2.25) and a standard microscope slide (0.15 c) makes the AG3D toolkit highly affordable even if one considers the addition of platinum electrodes. While the system described here used research grade powerpacks (>$500), electrophoresis could easily be achieved using 5 standard 9V batteries connected in series resulting in a further reduction in cost allowing for application in low-resourced research and educational environments. When coupled with affordable DIY detection systems [14] the AG3D allows for increased access to a cornerstone molecular biology technique.

## Ethics statements CRediT author statement

Conceptualisation: Garth LAbrahams, Earl Prinsloo; Data curation: Matthew W Barwick, Sidne Fanucci, Earl Prinsloo; Formal analysis: Matthew W Barwick, Sidne Fanucci, Khanyisile Buthelezi, Earl Prinsloo, Garth L Abrahams; Investigation: Sidne Fanucci, Matthew W Barwick; Methodology: Matthew W Barwick, Sidne Fanucci, Earl Prinsloo; Project administration: Earl Prinsloo, Garth L Abrahams; Resources: Earl Prinsloo, Garth L Abrahams; Design: Earl Prinsloo, Sidne Fanucci; Supervision: Earl Prinsloo, Garth Abrahams; Validation: Matthew W Barwick, Siso Mbanxa, Sidne Fanucci, Michael D Jukes; Visualisation: Siso Mbanxa, Matthew W Barwick; Writing – original draft: Earl Prinsloo, Sidne Fanucci, Matthew W Barwick, Khanyisile Buthelezi; Writing – review & editing: Earl Prinsloo, Sidne Fanucci, Matthew W Barwick, Khanyisile Buthelezi, Siso Mbanxa, Michael D Jukes, Garth L Abrahams.

## Acknowledgments

The authors acknowledge the National Research Foundation of South Africa (NRF UID: 129400), South African Medical Research Council (SAMRC), Council for Scientific & Industrial Research (CSIR) and Rhodes University for research and student funding. SF, SM are recipients of NRF bursary funding. KB is a recipient of DST-CSIR student funding. The authors further acknowledge Dr Ronen Fogel for constructive recommendations with respect to electrode selection and providing manuscript feedback.

## Declaration of interests

☒ The authors declare that they have no known competing financial interests or personal relationships that could have appeared to influence the work reported in this paper.

□ The authors declare the following financial interests/personal relationships which may be considered as potential competing interests:

## Notes

https://osf.io/fy7ca/

### Competing Interest Statement

The authors have declared no competing interest.

